# Chemogenetic Inhibition of Corticostriatal Circuits Reduces Cued Reinstatement of Methamphetamine Seeking

**DOI:** 10.1101/2021.07.28.453847

**Authors:** Angela M. Kearns, Benjamin M. Siemsen, Jordan L. Hopkins, Rachel A. Weber, Michael D. Scofield, Jamie Peters, Carmela M. Reichel

**Affiliations:** Department of Neuroscience, Medical University of South Carolina, Charleston SC 29425; Department of Anesthesiology, Medical University of South Carolina, Charleston SC 29425; Department of Anesthesiology, University of Colorado Denver, Anschutz Medical Campus, Aurora, CO 80045; Department of Pharmacology, University of Colorado Denver, Anschutz Medical Campus, Aurora, CO 80045

## Abstract

Methamphetamine (meth) causes enduring changes within the medial prefrontal cortex (mPFC) and the nucleus accumbens (NA). Projections from the mPFC to the NA have a distinct dorsal-ventral distribution, with the prelimbic (PL) mPFC projecting to the NAcore, and the infralimbic (IL) mPFC projecting to the NAshell. Inhibition of these circuits has opposing effects on cocaine relapse. Inhibition of PL-NAcore reduces cued reinstatement of cocaine seeking and IL-NAshell inhibition reinstates cocaine seeking. Meth, however, exhibits a different profile, as pharmacological inhibition of either the PL or IL decrease cued reinstatement of meth-seeking. The potentially opposing roles of the PL-NAcore and IL-NAshell projections remain to be explored in the context of cued meth seeking. Here we used an intersectional viral vector approach that employs a retrograde delivery of Cre from the NA and Cre-dependent expression of DREADD in the mPFC, in both male and female rats to inhibit or activate these parallel pathways. Inhibition of the PL-NAcore circuit reduced cued reinstatement of meth seeking under short and long-access meth self-administration and after withdrawal with and without extinction. Inhibition of the IL-NAshell also decreased meth cued reinstatement. Activation of the parallel circuits was without an effect. These studies show that inhibition of the PL-NAcore or the IL-NAshell circuits can inhibit reinstated meth seeking. Thus, the neural circuitry mediating cued reinstatement of meth seeking is similar to cocaine in the dorsal, but not ventral, mPFC-NA circuit.

## 1. Introduction

According to the United Nations Office on Drugs and Crime (2017), methamphetamine (meth) and other amphetamine-like stimulants have a pronounced burden of disease risk. Meth has long-term physiological and cognitive consequences including deficits in episodic memory, motor and language skills, executive function, and emotional regulation^1,2^. Many of the pathologies are driven by meth-induced changes in cortical neural plasticity impacting both top-down and bottom-up processing^3,4^; interfering with processing sensory information as well as cognition.

The medial prefrontal cortex (mPFC) is one of the main structures regulating relapse^5^ with glutamatergic corticostriatal projections at the forefront of addiction neurocircuitry^6,7^. Meth causes enduring changes within the mPFC including augmented burst firing within glutamatergic neurons^8^, changes in extracellular glutamate concentrations^8,9^, increased NMDA receptor expression, and adaptations in pre and post synaptic glutamate transmission^9–11^. Anatomically, projections from the mPFC to the nucleus accumbens (NA) have a distinct dorsal-ventral anatomical distribution^12^. The vast majority of neurons originating in the prelimbic (PL, i.e., the dorsomedial) cortex preferentially project to the NAcore, whereas the majority of neurons in the infralimbic (IL, i.e., the ventromedial) cortex project to the NAshell. These distinct anatomical projections serve distinct functional roles in cocaine relapse^13^. PL-NAcore inhibition reduces cued reinstatement of cocaine seeking following extinction^13–15^, whereas IL-NAshell inhibition reinstates previously extinguished rats^13^. Conversely, activation of the IL-NAshell circuit reduces cued reinstatement of cocaine seeking in extinguished rats^16,17^.

The projection neurons in the mPFC can act as promoters or inhibitors of drug seeking depending on if their downstream target is the NAcore or NAshell, respectively^18^. Interestingly, this dichotomy may be exclusive to cocaine as inhibition of the PL with Baclofen/Muscimol reduces both cocaine and heroin seeking; whereas the same manipulation of the IL promotes cocaine seeking and inhibits heroin seeking^19^. However, the existence of this duality within the stimulant drug class and specifically within meth relapse, has not been directly evaluated. As described above, we suspect that circuitry recruited to mediate reinstated meth seeking may be partially distinct from cocaine, as pharmacological inhibition of either mPFC subregion decreases cued reinstatement of meth seeking^20^, akin to heroin^21^. Downstream from the mPFC, pharmacological inhibition of the NAcore, but not NAshell, decreases cued reinstatement of meth seeking^20^. As such, the functional roles of the PL-NAcore and IL-NAshell remain unclear with respect to meth seeking, and all of the aforementioned studies were conducted exclusively in male subjects. Here we set out to determine the role of the mPFC-NA dorsal and ventral subcircuits in reinstated meth seeking triggered by conditioned drug cues using a Cre-dependent designer receptor exclusively activated by designer drugs (DREADD)-based approach (see Fig 1A schematic) in both male and female rats.

**Figure 1.**
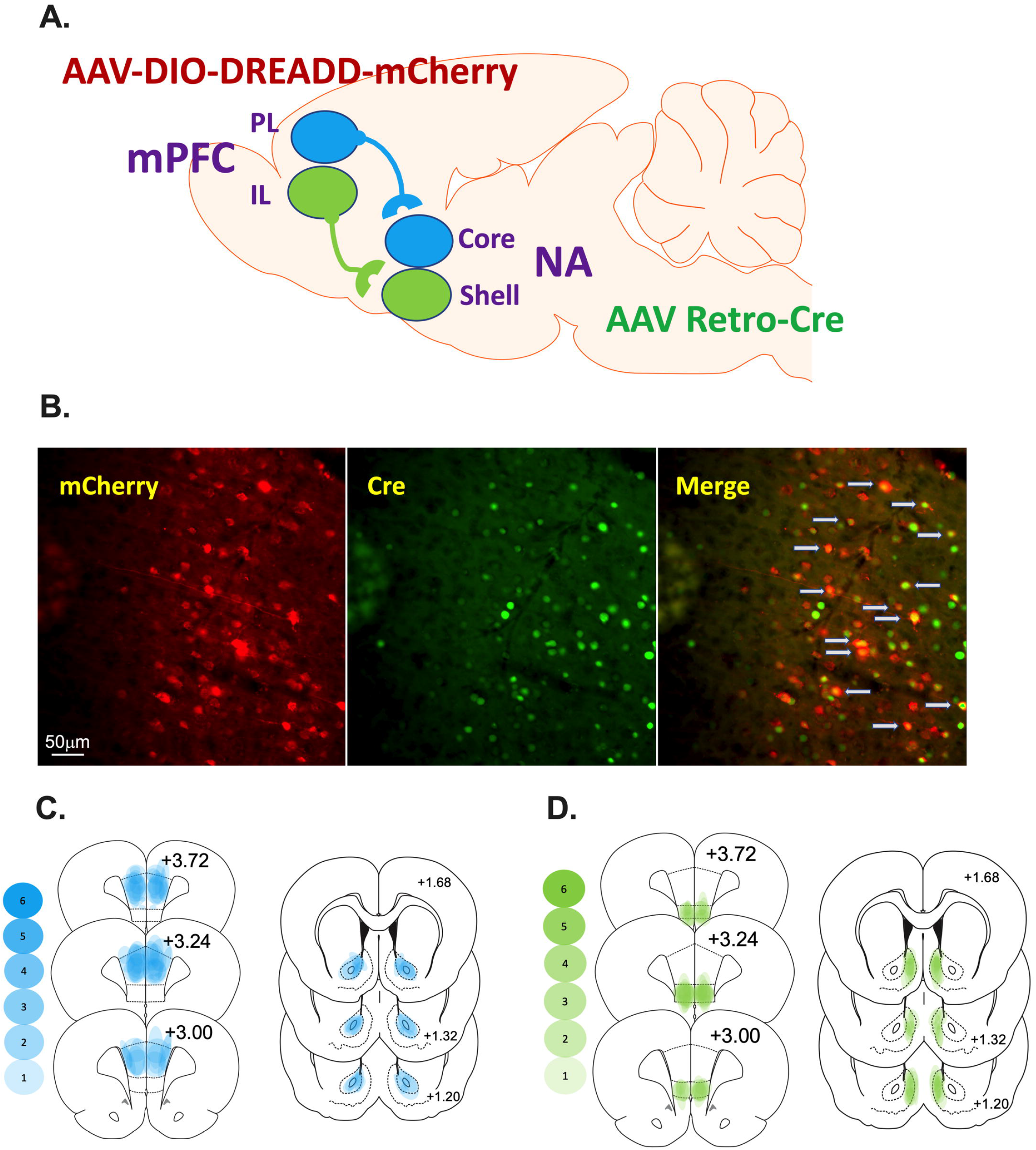
Schematic representation of the prelimbic (PL)-nucleus accumbens (NA)core and infralimbic (IL)-NAshell subcircuits as well as the dual viral approach. **A.** AAVrg-pmSyn1-enhanced blue fluorescent protein(eBFP)-Cre or AAVrg-hSyn-HI-enhanced green fluorescent protein(eGFP)-Cre was microinjected into the NAcore or NAshell depending on the experiment. The Cre-dependent Gi-DREADD [AAV-hSyn-DIO-hM4D(Gi)-mCherry or Gq-DREADD [AAV-hSyn-DIO-hM3D(Gq)-mCherry] into the PL or IL (i.e., the cell body area) depending on the experiment. As a control, some rats received DIO-mCherry [AAV-hSyn-DIO-mCherry] instead of DREADDs into the cortical area under similar parameters. **B.** (Left panel) IHC detection of DIO-hM4Di-mCherry (red) in the PL. (Center panel) IHC detection of GFP (green) in the PL. (Left panel). Overlay of the cells expressing Cre merged with those expressing mCherry are identified with gray arrows (Right panel). **C and D.** Heat maps were generated for the PL-NAcore (C) and IL-NAshell (D) to represent the extent of viral transduction in each region. Transparent blue or green shapes were used to yield more saturation when viral placements were consistent across animals.

## 2. Materials & Methods

### 2.1 Subjects

Age-matched male (250-275g) and female (225-250g) Sprague Dawley rats (Envigo, Indianapolis, IN, USA) were used in these experiments. Details are supplied in the <u>Supplemental Information</u>.

### 2.2 Surgery

Catheter surgeries are described in the <u>Supplemental Information</u>.

### 2.3 Virus microinjections

Figure 1A is a schematic representation of the dual viral approach and the PL-NAcore and IL-NAshell subcircuits. Microinjections of the viral constructs were infused under the same anesthetic plane as catheter surgery. AAVrg-pmSyn1-enhanced blue fluorescent protein(eBFP)-Cre [5×10^12^ vg/ml; Addgene #51507] or AAVrg-hSyn-HI-enhanced green fluorescent protein(eGFP)-Cre-WPRE-SV40 [7×10^12^ vg/ml; Addgene #105540]) was microinjected into the NAcore or NAshell (0.5ul/side at 0.15ul/min). The Cre-dependent DREADDs (Gi-DREADD: AAV2-hSyn-DIO-hM4D(Gi)-mCherry Addgene #44362 or Gq-DREADD: AAV2-hSyn-DIO-hM3D(Gq)-mCherry Addgene #44361; 0.6 μL/side at 0.15μL/min; 5×10^12^ vg/ml) were injected into the PL or IL. As a control, some rats received AAV2-hSyn-DIO-mCherry Addgene #50459 (0.6 μL/side at 0.15μL/min) instead of DIO-DREADDs.

During intracranial surgery, the virus was infused via Nanoject II at a volume of 50.6 nL per injection every 30 seconds at a rate of 23 nL/second followed by an additional 10 minutes to allow the injected virus to diffuse prior to removal of the pipette. Stereotaxic coordinates for the PL were AP +2.8 mm; ML, ± 0.64 mm; DV −3.7 mm, and for the NAcore: AP +1.6 mm; ML ± 2.8 mm (10° angle); DV −7.1 mm. Stereotaxic coordinates for the IL were AP + 3.0 mm; ML, ± 0.64 mm; DV −5.5 mm, and for the NAshell: AP +1.7 mm; ML, ± 0.8 mm, DV −7.0 mm. Figure 1B shows representative images of the DIO-hM4Di-mCherry in the PL, green fluorescent protein (GFP) tag on the AAVrg-Cre in the cell body locus demonstrating effective retrograde transport of the construct, and the overlay between the mCherry tag on the DIO-hM4Di DREADD construct (red) and IHC detection of GFP (green) overlay demonstrating that the cells expressing Cre are those that project to the NAcore. Representative images of the GFP expression in the terminal areas, NAcore and NAshell, are depicted in Supplemental Figure 1. Heat maps were generated for the PL-NAcore (Figure 1C) and IL-NAshell (Figure 1D) to represent the extent of mCherry/GFP overlap in the cortex and GFP in the NA.

### 2.4 Meth or Sucrose Self-Administration

In all experiments, except one, rats self-administered methamphetamine hydrochloride (Sigma, St Louis, MO). The exception is when sucrose was used as the reinforcer. Chamber and procedural details are described in the <u>Supplemental Information</u>.

### 2.5 Extinction, Abstinence, and Cued Reinstatement Testing

Extinction and abstinence are described in the <u>Supplemental Information</u>. On the cued reinstatement test session, rats were pretreated with vehicle (5% dimethylsulfoxide (DMSO) + 95% saline) or Clozapine-N-oxide (CNO, 3 or 10 mg/kg) 30 min before chamber placement. CNO was dissolved in 5% dimethylsulfoxide (DMSO) + 95% saline and administered at a dose of 10 and 3 mg/kg i.p. based on reports from our lab and others that this may be the minimal effective dose for use in this system^16,22–25^. All rats tested with vehicle had 1-2 additional cue tests with 3 or 10 mg/kg CNO. Each animal has an extinction value (mean of last 2 extinction sessions). Each rat tested on vehicle. The vehicle test was counter-balanced with the CNO tests such that an equal number of animals received vehicle or CNO on the first test. When rats were tested on both 3 and 10 mg/kg CNO the 3 test conditions (veh, 3, and 10 mg/kg CNO) were counter-balanced. Supplemental Figure 2 details when these tests occurred. Rats were kept at 2-3 cue tests in order to limit potential test ordering effects resulting from within trial extinction of the reward-associated cues. Test sessions were 1-2 hours depending on if rats were abstinent or reinstated, respectively. During the cue test session, a response on the active lever resulted in the presentation of the light + tone stimulus previously paired with reward (meth or sucrose) delivery; however, no reward was delivered.

### 2.6 Immunohistochemistry

At the end of the experiments, rats were transcardially perfused and brains removed. Tissue was sliced on a cryostat (40 um) and underwent immunohistochemical procedures before being mounted onto slides, dehydrated, cover- slipped, and examined under a microscope to visualize viral expression. Subjects were eliminated from the final dataset if no expression was visible or if there was spread of transduced cells outside of the target region and into adjacent regions. This analysis was performed using cortical subregion boundaries defined by coronal slices from the atlas of Paxinos and Watson (2007). Antibodies are detailed in the <u>Supplemental Information.</u>

### 2.7 Confocal and Microscopy and Analysis

All imaging and analysis were performed by an investigator blind to experimental groups. Fos and mCherry co-localization imaging and analysis was performed as previously described^26^. Additional details for the confocal and microscopy and analysis are in the <u>Supplemental Information.</u>

### 2.8 Validation of DIO-hM4Di and DIO-hM3Dq viral vectors

To validate the function of CNO on DIO-hM4Di, a subset of animals had DIO-hM4Di infused into the PL and AAVrg-Cre in the NAcore. These rats underwent meth self-administration (described in Supplemental Information). Rats were then re-exposed to the meth associated cues 30 min after administration of 10 mg/kg CNO or vehicle. It is important to note that these rats underwent cue exposure to induce Fos expression to test inhibition. The test session lasted 2hr at which time rats were perfused and brain tissue processed for detection of mCherry and Fos immunoreactivity. To validate the function of CNO on DIO-hM3Dq, a separate group of rats had DIO-hM3Dq infused into IL and AAVrg-Cre into the NAshell. For both hM4Di and hM3Dq data sets, images were collected and analyzed to determine numbers of Fos+ and mCherry+ cells. Within our analyses data are presented as the number of double labeled mCherry+Fos+ cells normalized to total mCherry cells per brain section as described above.

### 2.9 Determining the role of the PL-NAcore circuit during cued meth seeking

We conducted a series of experiments to evaluate the PL-NAcore circuit in cued reinstatement to meth seeking. Previous data have implicated the PL area of the mPFC as an important substrate in reinstated meth seeking^20,27^ and the PL neurons projecting to the NAcore are crucial for cued reinstatement of cocaine seeking^15,28^. In all experiments, rats underwent surgery, meth self-administration and were then subsequently tested for lever pressing following abstinence (with or without extinction). The timeline and individual experiments are depicted in Supplemental Figure 2. In Experiments 1 and 2, AAVrg-Cre was infused into the NAcore in order to retrogradely deliver the Cre-recombinase gene to the PL. The Cre-dependent DREADD virus AAV-DIO-hM4Di was infused in the PL to allow for selective expression of inhibitory DREADD in PL-NAcore neurons (see Schematic Fig 1). In Experiment 1, rats were tested for cued reinstatement following extinction and in Experiment 2 rats were tested for lever responding after abstinence. In Experiment 3, AAVrg-Cre was infused into the NAcore and the Cre-dependent DREADD virus AAV-DIO-hM3Dq into the PL and rats were tested for cued reinstatement of meth seeking. In Experiment 4, AAVrg-Cre was infused into the NAcore, but the PL was infused with Cre-dependent mCherry (DIO-mCherry) that lacked the DREADD and rats were tested for cued reinstatement of meth seeking.

### 2.10 Evaluation of the PL-NAcore circuit on cued meth seeking following long assess meth or sucrose self-administration

To determine if DIO-hM4Di inhibition of the PL-NAcore circuit extends to a long-access model of meth “addiction” or generalizes to non-drug reward, a subset of male rats self-administered meth or sucrose for longer durations (8-hr) in daily sessions for 15 days on an FR1 schedule of reinforcement. In Experiment 5, AAVrg-Cre was infused into the NAcore and the Cre-dependent DREADD virus AAV-DIO-hM4Di was infused in the PL; rats tested for reinstatement in response to meth associated cues. In parallel, Experiment 6 was identical except the reinforcement was sucrose rather than meth.

### 2.11 Evaluation of the PL-NAcore circuit on cued meth seeking

Previously, temporary inactivation of the NAshell with the GABA agonists baclofen and muscimol (B/M) did not alter cued or primed reinstatement of meth seeking, yet inactivation of the IL specifically reduced the cued reinstatement of meth seeking^20^. Therefore, to clearly define the role of this projection, we inhibited and activated this projection in separate experiments. In Experiment 7, AAVrg-Cre was infused into the NAshell to retrogradely deliver Cre-recombinase and the IL was infused with Cre-dependent inhibitory DREADD (DIO-hM4Di). In Experiment 8, AAVrg-Cre was infused into the NAshell and the IL was infused with AAV-DIO-hM3Dq to activate the circuit. In both experiments rats tested for cued reinstatement of meth seeking.

### 2.9 Experimental Design and Data Analysis

Both sexes are included in these studies because differences occur in metabolism^29^ and acquisition of drug self-administration^30,31^, and in basic and synaptic excitatory transmission^11,32^. However, our behavioral endpoint, cued reinstatement, is not sensitive to sex as a biological variable^33,34^ unless the response ratio is increased during the reinstatement test^33^. Because of this, we predict that the PL-NAcore and the IL-NAshell will mediate cued reinstatement when data is collapsed across both sexes.

Active and inactive lever responding from the self-administration phase were analyzed with a 3-way mixed analysis of variance (ANOVA) with sex (male and female), lever (active and inactive), and day (1-15) as the independent variables. The number of infusions received and extinction data were analyzed with a 2-way mixed ANOVA with sex and day as the independent variables. Data is presented collapsed across sex due to lack of significant sex differences. The data are disaggregated by sex in the supplemental data. Also, due to very limited responding on the inactive lever on test day, the means and SEMs are presented in Supplemental Table 1. Reinstatement data were analyzed with repeated measures or a Mixed-effects analysis to account for missing data because not all rats tested on each reinstatement test (See Supplemental Figure 1). Fos data were analyzed with an unpaired two-tailed t-test. Post-hoc comparisons were conducted with Holm-Sidak’s multiple comparisons test to control for family-wise error. Statistical significance was set at an alpha of p<0.05.

## 3. Results

### 3.1 CNO functionally inhibits or activates Fos expression

To validate the function of CNO on DIO-hM4Di, animals (n=8) had DIO-hM4Di infused into the PL and AAVrg-Cre in the NAcore. Tissue from two rats was damaged and not used in the analysis. As expected, CNO reduced Fos expression in mCherry+ cells in the PL [Fig 2A; t(4)=3.35, p=0.03]. While we specifically focused on Fos expression within virally transduced cells, the number of mCherry+ cells in the PL did not differ between vehicle (5% DMSO) or CNO treatment groups [t(4)=0.32, p=0.764]. To validate the function of CNO on DIO-hM3Dq, a separate group of rats had DIO-hM3Dq infused into IL and AAVrg-Cre into the NAshell. As expected, an injection of CNO-induced Fos expression in mCherry+ cells in the IL relative to vehicle [Fig 2B; t(7)=11.01, p=0001]. Here as above, the number of mCherry+ cells in the IL did not differ between rats that received vehicle (5% DMSO) or CNO injection [t(7)=1.68, p=0.14]. These experiments were designed to validate the efficacy of our DIO-hM4Di and DIO-hM3Dq vectors. Additional information regarding the regional specificity of DREADD-mCherry+ expression and the extent and localization of Fos induction is depicted in Supplemental Figure 3.

**Figure 2.**
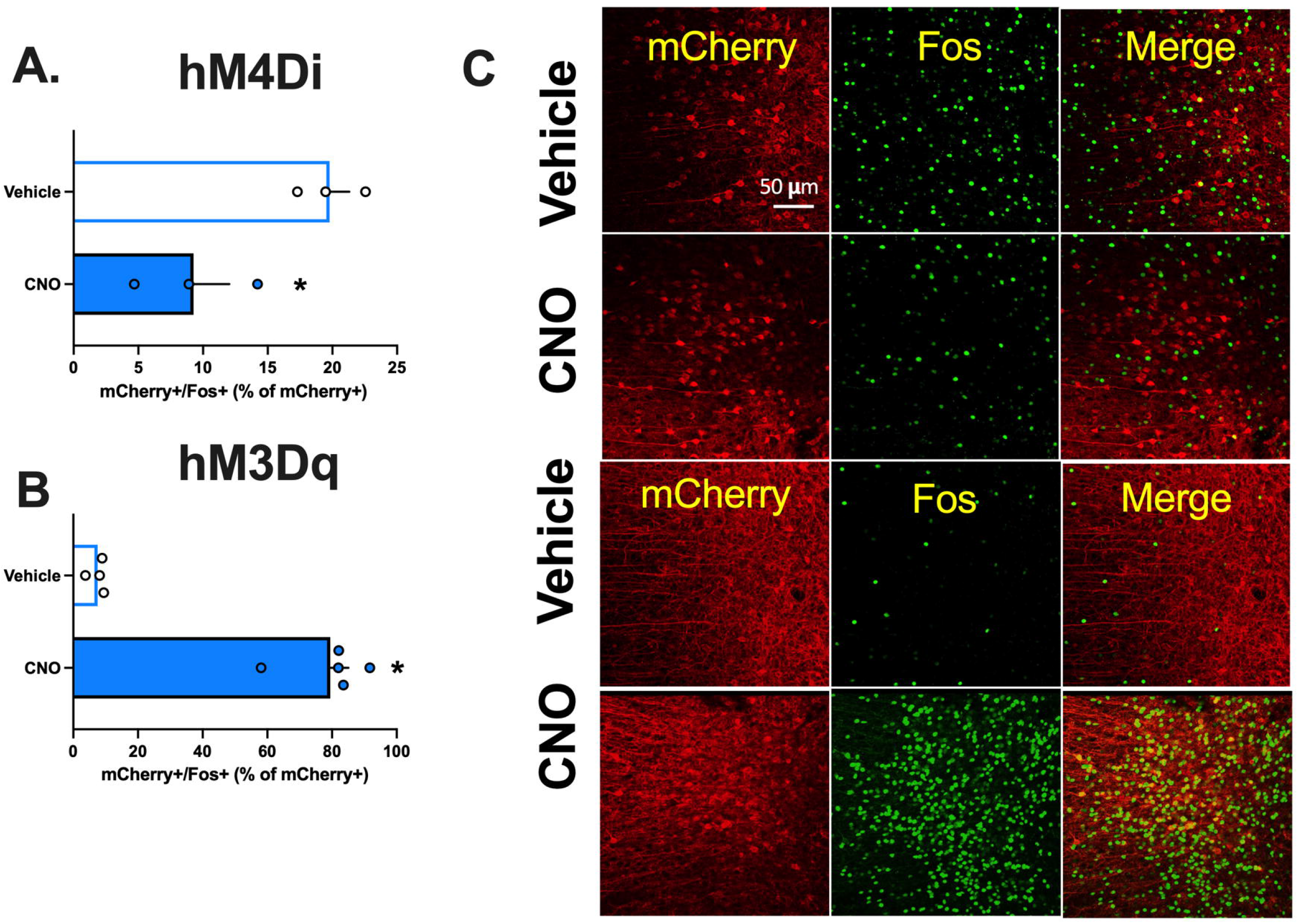
Validation of clozapine-N-oxide’s (CNO) function on DREADD activity and inhibition. **A.** CNO reduced mCherry+/Fos+ cells in the infralimbic (IL) cortical projecting neurons to the nucleus accumbens (NA)shell during cued meth seeking **B.** CNO increased mCherry+/Fos+ cells in the prelimbic (PL) cortical projecting neurons to the nucleus accumbens (NA) core. **C.** Representative images depicting the co-labeling of mCherry positive (marker of DREADD) and Fos positive neurons (shown here in green). *Significant difference from VEHICLE.

### 3.2 Experiment 1. Inactivation of the PL projections to the NAcore reduces cued reinstatement of methamphetamine seeking

For Experiments 1–4, which focus on PL-NAcore inhibition during cue-induced reinstatement of meth seeking, experimental timelines are depicted in Figure 3A. Given that self-administration and extinction lever presses and infusions earned were consistent across experiments, these data were collapsed across experiments for self-administration (Fig 3B) and extinction (Fig 3C) (See Supplemental Figure 4 for separate graphs and analysis for each experiment).

**Figure 3.**
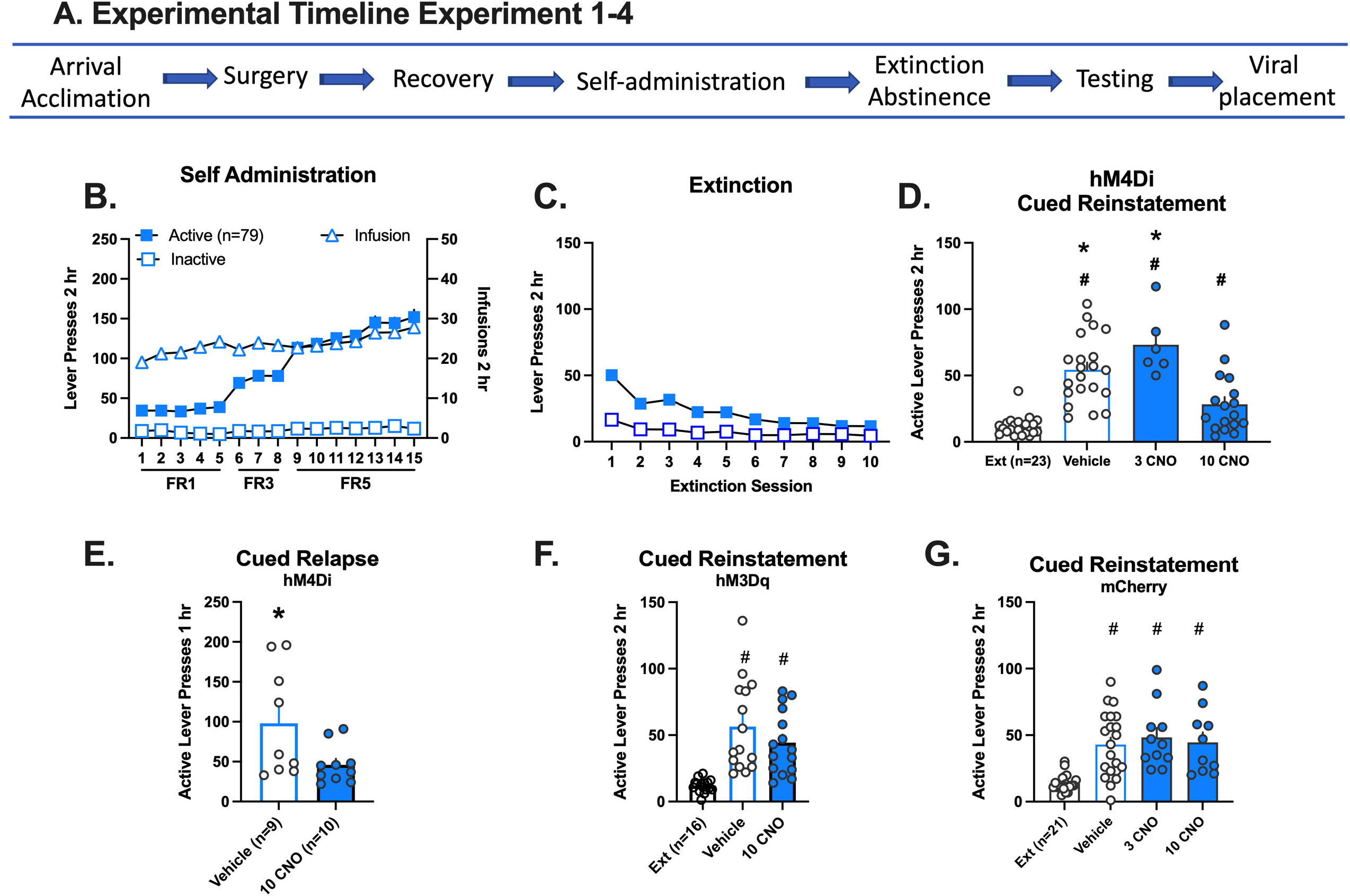
Inhibition and activation of the prelimbic (PL)-nucleus accumbens (NA)core circuit and methamphetamine seeking in response to drug cues. **A.** Experimental timeline for experiments 1–4. **B.** Active and inactive lever responding (left axis) and meth infusions (right axis) over 15 daily 2 hr sessions. Self-administration data is combined for all experiments represented in D-G. The individual graphs per experiment are disaggregated by sex and depicted in Supplement Figure 4. **C.** Active and inactive lever responding over 10 daily extinction (Ext) sessions. Extinction data is combined for all experiments represented in D-G. The individual graphs per experiment are disaggregated by sex and depicted in Supplemental Figure 4. **D.** Active lever presses during extinction (calculated as the mean of the last 2 extinction sessions) and in response to presentation of meth associated cues following an injection of vehicle, 3 or 10 mg/kg CNO. All groups increased active lever presses relative to extinction in rats with DIO-hM4Di in the PL. However, lever responding following 10 mg/kg clozapine-N-oxide (CNO) was significantly decreased relative to vehicle and 3 mg/kg CNO. **E.** Active lever presses following abstinence in response to an injection of vehicle or 10 mg/kg CNO. In rats with DIO-hM4Di in the PL, lever responding decreased following an injection of 10 mg/kg CNO relative to VEHICLE following re-exposure to the meth context after abstinence. **F.** Active lever presses during extinction and in response to presentation of meth associated cues following an injection of vehicle or 10 mg/kg CNO in rats with DIO hM3Dq in the PL. Both test conditions increased responding relative to extinction. **G.** Active lever presses during extinction and in response to presentation of meth associated cues following an injection of vehicle, 3 or 10 mg/kg CNO. Rats with DIO-mCherry in the PL rather than the DREADD increased lever responding relative to extinction across all test conditions. (G) The schematic represents the extent of mCherry expression in the anteroposterior representation of the prefrontal cortex. *Significant difference from 10 mg/kg CNO. #Significant difference from extinction.

During self-administration (Fig Supplemental Figure 4A), active lever pressing increased over days in males (n=11) and females (n=12), which reflects the change in FR ratio [lever × day interaction, F(14,672)=68.87, p=0.0001]. The number of infusions earned stayed consistent across the 15 days. During extinction, lever responding decreased over days [sex × day interaction, F(9,202)=2.8, p=0.004, Fig Supplemental Figure 4B]; post hoc comparisons between males and females did not reach statistical significance. On the cued reinstatement test, 10 mg/kg CNO (injected 30 min before the session) significantly reduced lever pressing relative to vehicle [Fig 3D, main effect of treatment, F(3,39)=25.73, p=0.0001]. Post-hoc comparisons confirm that all groups reinstated relative to extinction (vehicle and 3 CNO Holm-Sidak’s, p<0.0001; 10 CNO Holm-Sidak’s, p=0.012), but 10 mg/kg CNO were significantly lower than vehicle or 3 mg/kg CNO (p’s=0.008). No sex differences were detected (Supplemental Figure 4C). Five rats were excluded from the analysis: One rat failed to extinguish responding, two had missed placements, and two died during surgery.

### 3.3 Experiment 2. Inactivation of the PL-NAcore reduces relapse after abstinence

To determine if reduced meth seeking following inhibition of the PL-NAcore projection requires extinction training, we used a short term, forced abstinence protocol^35^ that excludes extinction^36,37^. During self-administration (Supplemental Figure 4D) active lever pressing increased over days [males (n=10) and females (n=9), lever × day interaction, F(14,400)=16.46, p=0.0001] as did the number of infusions received [main effect of day, F(14,223)=5.72, p=0.0001]. Rats were tested once with CNO 10 mg/kg or VEHICLE under the same parameters as cued reinstatement but after a 7-day home-cage abstinence period. Again, under these conditions, CNO significantly decreased meth seeking on the active lever relative to VEHICLE [main effect of test, Fig 3E, F(15)=4.86p=0.043] in both sexes (Supplemental Figure 3E). There were no interactions or sex effects. Two males and one female did not wake from surgery, and 3 males were excluded for catheter blockage.

### 3.4 Experiment 3. Activation of the PL-NAcore does not induce reinstatement to meth seeking

To determine whether the PL projection to the NAcore is a driver of relapse, we infused AAVrg-Cre in the NAcore and the Cre-dependent excitatory DREADD (DIO-hM3Dq) in the PL. During self-administration (Supplemental Figure 4F), active lever pressing increased over days in male (n=8) and female (n=8) rats [lever × day interaction, F(14,391)=7.67, p=0.0001] as did the number of infusions received [main effect of day, F(14, 195)=2.85, p=0.0006]. During extinction (Supplemental Figure 4G), lever responding decreased over days [main effect of session, F(9,14)=9.77, p=0.0001]. During reinstatement testing (Fig 3F), rats reinstated regardless of treatment group [main effect of test, F(2,28)=19.34, p=0.0001] or sex [Supplemental Figure 4H].

### 3.5 Experiment 4. CNO does not impact reinstatement to meth seeking in DIO-mCherry rats

An “empty” virus control experiment confirmed that the above finding resulted from hM4Di inhibition of the PL-NAcore pathway and not administration of CNO. As before, the AAVrg-Cre was infused into the NAcore, but the PL was infused with Cre-dependent mCherry (DIO-mCherry) that lacked the DREADD. During self-administration (Supplemental Figure 4I), active lever pressing increased over days in both males (n=10) and females (n=11) [lever × day interaction, F(14,524)=52.6, p=0.0001]. The number of infusions received increased over days [main effect of day, F(14,262)=2.64, p=0.0013]. During extinction (Supplemental Figure 4J), lever responding decreased over days [main effect of session, F(9,171)=21.63, p=0.0001]. During reinstatement testing (Fig 3G), all groups increased lever pressing in response to the meth-associated cue [main effect of test, F(3,38)=13.22, p=0.0001, post hocs p<0.0001]. This experiment controls for any potential effects of 10mg/kg CNO. No sex differences were detected (Supplemental Figure 4K). When directly compared, CNO decreased lever responding in rats with hM4Di relative to DIO-mCherry (Supplemental Figure 5). These data demonstrate that neither viral transduction of the PL-NAcore pathway nor 10mg/kg CNO significantly impacts cued meth seeking.

### 3.6 Experiments 5 and 6. Inactivation of the PL projections to the NAcore selectively reduces cued reinstatement of methamphetamine seeking, but not sucrose seeking, under long-access conditions

To determine if DIO-hM4Di inhibition of the PL-NAcore circuit extends to a long-access model of meth “addiction” or generalizes to non-drug reward, a subset of male rats self-administered meth during longer duration (8-hr) daily sessions for 15 days on an FR1 schedule of reinforcement (Fig 4A). During acquisition, rats readily discriminated active from inactive lever over days [Fig 4B, Day × Lever interaction, F(14,232)=13.15, p=0.0001] and escalated drug intake on days 4-15 relative to day 1 as indexed by an increase in infusions over time [F(2,13)=29.63, p=0.0001, post hoc p<0.05]. Rats extinguished active lever responding over days [Fig 4C, Day × Lever interaction, F(7, 98)=3.64, p=0.0016]. On the cued reinstatement test (Fig 4D), both VEHICLE and CNO (10 mg/kg) treatment groups increased active lever responding relative to extinction, but inhibition of the PL-NAcore circuit with CNO again significantly reduced lever pressing relative to VEHICLE [main effect of test day, F(2,14)=20.20, p=0.0001, Holm-Sidak’s post hoc p<0.05]. These findings are limited to only having males in the experiment.

**Figure 4.**
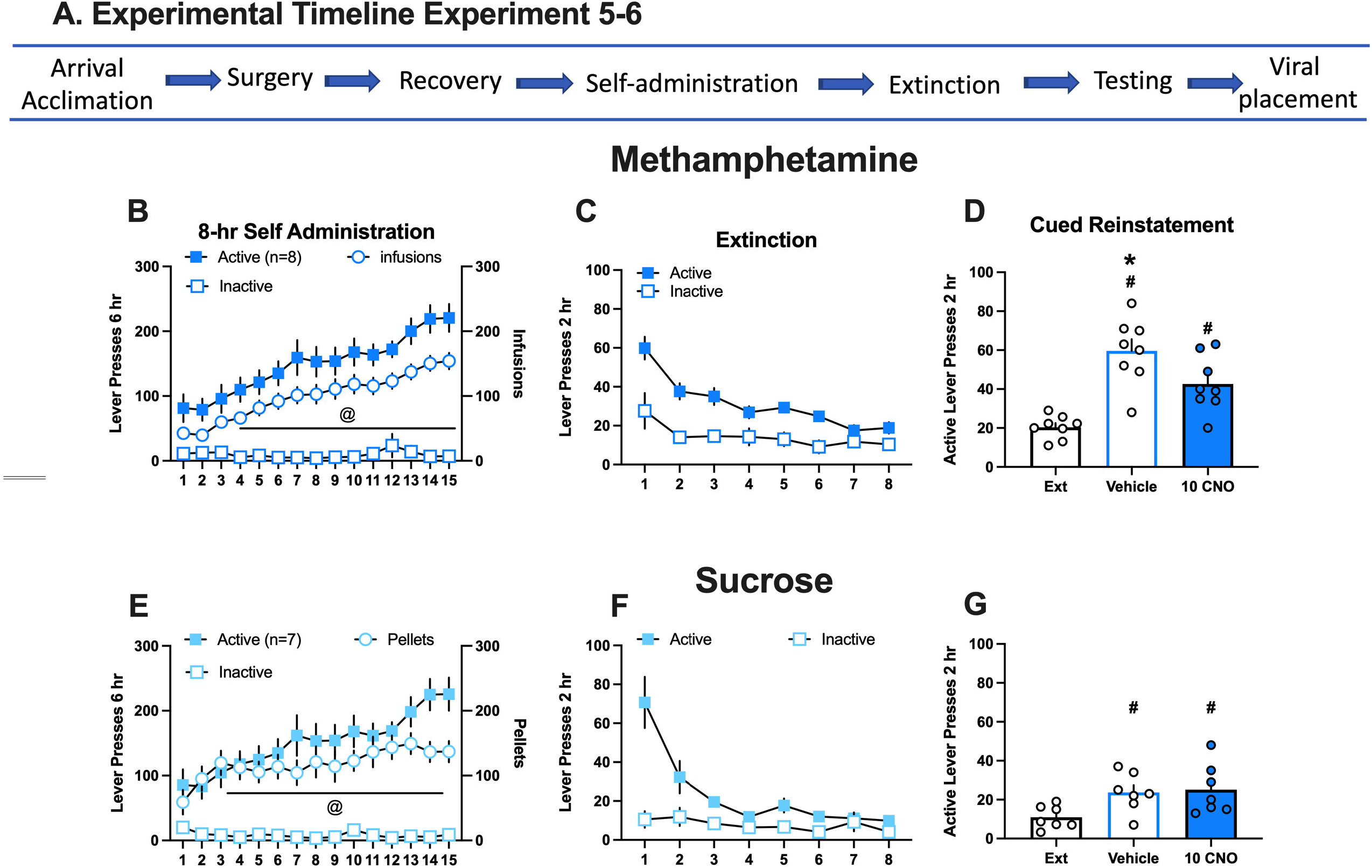
Inhibition of the prelimbic (PL)-nucleus accumbens (NA)core decreased methamphetamine seeking in male rats that underwent extended drug access but not sucrose. **A.** Experimental timeline for experiments 5 and 6. **B.** Active and inactive lever responding (left axis) and meth infusions (right axis) in rats over 15 daily 8 hr sessions. Active lever presses and infusions escalated over time. **C.** Active and inactive lever responding over 8 daily extinction sessions. Rats extinguished responding over time. **D.** Active lever presses during extinction (mean lever presses of the last 2 extinction sessions) and in response to presentation of meth associated cues following an injection of vehicle or 10 mg/kg clozapine-N-oxide (CNO). Active lever responding increased after VEHICLE and CNO injections in response to meth associated cues relative to extinction, but lever pressing was significantly reduced following 10 mg/kg CNO. **E.** Active and inactive lever responding (left axis) and sucrose pellets (right axis) in rats over 15 daily 8 hr sessions. Sucrose pellets earned stabilized over the sessions. **F.** Active and inactive lever responding over 8 daily extinction sessions. Rats extinguished lever responding over time. **G.** Active lever presses on extinction and in response to presentation of meth associated cues following an injection of vehicle or 10 mg/kg CNO. Lever presses increased in response to sucrose associated cues following VEHICLE and CNO injections relative to extinction, but there were no differences between treatments. @Significant increase in reinforcer (meth or sucrose) over time. *Significant difference from 10 mg/kg CNO. #Significant difference from extinction.

Sucrose control rats that underwent similar long-access (8-hr daily) conditions also readily discriminated the active from inactive lever over days [Fig 4E, Day × Lever interaction, F(14,165)=10.56, p=0.0001] and increased sucrose pellets over time [main effect of day, F(4,22)=3.37, p=0.03]. Sucrose rats also extinguished active lever responding over days [Fig 4F, Day × Lever interaction, F(7, 84)=12.07, p=0.0001]. On the cued reinstatement test (Fig 4G), CNO had no impact on sucrose seeking when compared to VEHICLE, indicated by active lever responding relative to extinction baseline [main effect of test day, F(2,12)=6.57, p=0.0118, post hoc p<0.05], as no effect of treatment or interaction was observed. Taken together, these data demonstrate that PL neurons projecting to the NAcore have a critical role in cued reinstatement of meth, but not sucrose, seeking following extinction. These data also indicate that neither 10mg/kg CNO nor DIO-hM4Di inhibition of the PL-NAcore circuit have a non-specific effect on reward seeking.

### 3.7 Experiment 7. Inhibition of the IL projections to the NAshell reduced cued reinstatement of methamphetamine seeking following extinction

Previously, temporary inactivation of the NAshell with the GABA agonists baclofen and muscimol (B/M) did not alter cued or primed reinstatement of meth seeking, yet inactivation of the IL specifically reduced the cued reinstatement of meth seeking^20^. Therefore, to clearly define the role of this projection, we inhibited or activated this projection in separate experiments. The general experimental timeline is depicted in Figure 5A. Here as above, data were combined across experiments for self-administration and extinction in Figures 5B and 5C, respectively. In Experiment 7, AAVrg-Cre was infused into the NAshell to retrogradely deliver Cre-recombinase and the IL was infused with Cre-dependent inhibitory DREADD (DIO-hM4Di). During self-administration (Supplemental Figure 6A), active lever pressing increased over days in both male (n=6) and female (n=6) rats, [lever × day interaction, F(14, 319)=26.17, p=0.0001]. The number of infusions earned stayed consistent across the 15 days. During extinction (Supplemental Figure 6B) lever responding decreased over days [main effect of day, F(9,99)=21.02, p=0.0001]. On the cued reinstatement test (Fig 5D), rats reinstated regardless of treatment [main effect of test, F(3,30)=13.01, p=0.0001] or sex (Supplemental Figure 6C). CNO (10 mg/kg) inhibition of the IL-NAshell circuit significantly decreased reinstated responding relative to VEHICLE [Holm-Sidak’s post hoc, p<0.05]. Two female rats did not wake up from surgery and two male rats were excluded from the analysis for viral spread into adjacent regions.

**Figure 5.**
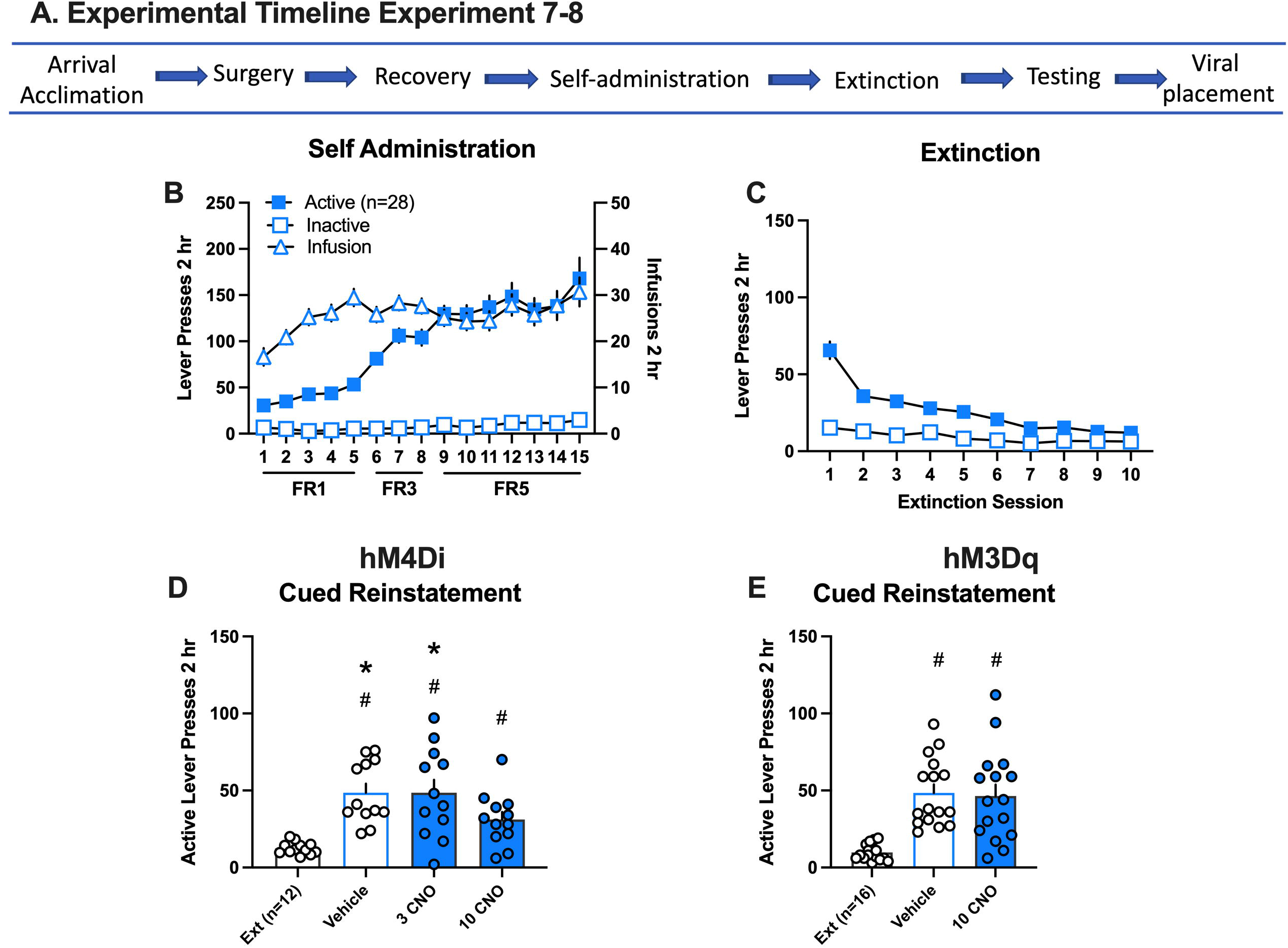
Inhibition and activation of the infralimbic (IL)-nucleus accumbens (NA)shell circuit and methamphetamine seeking in response to drug cues. **A.** Experimental timeline for experiments 7 and 8. **B.** Active and inactive lever responding (left axis) and meth infusions (right axis) over 15 daily 2 hr sessions. Self-administration data is combined for experiments represented in D and E. The individual graphs per experiment are disaggregated by sex and depicted in Supplemental Figure 5. **C.** Active and inactive lever responding over 10 daily extinction (Ext) sessions. Extinction data is combined for all experiments represented in D and E. The individual graphs per experiment are disaggregated by sex and depicted in Supplemental Figure 5. **D.** All groups increased active lever presses relative to extinction in rats with DIO-hM4Di in the IL. However, lever responding following 10 mg/kg CNO was significantly decreased relative to vehicle and 3 CNO tests. **D.** Reinstatement levels were similar in rats with DIO-hM3Dq in the IL in response to VEHICLE or CNO. *Significant difference from 10 mg/kg CNO. #Significant difference from extinction.

Given that activation of the IL-NAshell reduces reinstatement of cocaine seeking in response to cocaine associated cues, we sought to investigate the role of this projection in cued meth seeking. In Experiment 8, we infused AAVrg-Cre in the NAshell and the Cre-dependent excitatory DREADD (DIO-hM3Dq) in the IL. During self-administration (Supplemental Figure 6D), active lever pressing increased over days in males (n=8) and females (N=9), [lever × day interaction, F(14,392)=18.15, p=0.0001]. The number of infusions received increased over days [main effect of day, F(14,196)=4.82, p=0.0001]. During extinction (Supplemental Figure 6E), lever responding decreased over the days [main effect of session, F(9,117)=20.78, p=0.0001]. During reinstatement testing (Fig 5E), rats reinstated lever pressing in response to the meth-associated cues regardless of treatment [main effect of test, F(2,28)=19.65, p=0.0001, post hocs p<0.05] or sex (Supplemental Figure 5F), and there was no significant difference in reinstatement between treatments.

## 4. Discussion

Activity in the PL promotes both cocaine and heroin seeking, whereas activity in the IL inhibits cocaine but promotes heroin seeking^19^. Until now, the functional roles of the PL-NAcore and IL-NAshell subcircuits were not defined for meth seeking. We demonstrate that activity in the PL-NAcore promotes cued reinstatement of meth seeking following extinction or forced abstinence and after long-access meth self-administration. Interestingly, activity in the IL-NAshell also promotes cued reinstatement of meth seeking after extinction, yet boosting activity in the IL-NAshell or PL-NAcore pathways has no additional impact on reinstatement, perhaps due to a ceiling effect in behaving animals. The pathway specific manipulations described do not result from non-specific effects of CNO given that mCherry expression without a DREADD had no impact on meth seeking.

### Role of the PL-NAcore in meth seeking

Consistent with the literature on cocaine^13,15,38^ and heroin^21^ the PL appears to function to promote meth seeking^20,27^. Pharmacological inhibition of PL reduces both cued and primed forms of meth reinstatement^20,39^, and optogenetic inhibition of PL also decreases meth-primed reinstatement^27^. To determine whether the stimulatory function of the PL is mediated via its projections to the NAcore, we chemogenetically inhibited activity in the PL-NAcore projection following extinction or abstinence from meth self-administration, including long-access conditions. Under all conditions tested, chemogenetic inhibition of the PL-NAcore circuit reduced meth seeking. These findings are consistent with previous observations that pharmacological inhibition of the NAcore decreases cued and primed reinstatement of meth seeking after extinction^20,39^ and voluntary abstinence^59^. We extend these findings to demonstrate subcircuit-specific roles within the mPFC-NA projections. Drug self-administration dysregulates glutamatergic signaling in the PL-NAcore. Specifically, PL neurons are thought to be hypoactive under basal conditions, a state described as “hypofrontality.” However, this may render them hyperexcitable by drug-associated stimuli, i.e., cues. In the presence of drug cues, PL neurons thus become active, resulting in enhanced glutamate release in the NAcore^40–42^. Normalization of glutamatergic signaling in this circuit alleviates drug seeking^5,43^. Thus, our findings support chemogenetic inhibition of the PL-NAcore circuit as a therapeutic strategy for normalizing activity in this circuit to reduce meth relapse. However, activation of this circuit did not exacerbate meth reinstatement as may have been expected, perhaps activation of the PL-NAcore neurons is already at a ceiling effect required to engage the behavior in reinstating animals.

We show here that PL-NAcore inhibition also decreased meth seeking following extended access to meth, but not sucrose. Longer self-administration sessions are considered more suitable for studying addiction processes in rodent models because increasing the session length results in an escalation of drug intake over time reminiscent of human compulsive drug taking^44^. Moreover, long-access meth enhances drug seeking and cognitive impairments (reviewed in ^2^) and results in persistent cortical alterations in monoamine systems^45^ and glutamatergic function^8,10^. Here we used a 6-hour sucrose self-administration protocol to match the rate of lever responding between meth and sucrose rats. Whereas meth rats escalated meth intake over time, sucrose rats did not. Instead, sucrose rats stabilized sucrose intake over the 15-day self-administration period and extinguished behavior rapidly. Pharmacological inhibition of the PL does not impact sucrose reinstatement^46,47^. Our findings corroborate this earlier work and extend our knowledge to a specific PL output, namely the PL-NAcore circuit. The purpose of sucrose animals in this study was to determine if PL-NAcore inhibition impacted a “natural” reward process. These data do not rule out the role of the PL-NAcore in pathological overeating because some forms of obesity are compulsive in nature and therefore consistent with an addictive disorder^48^.

### Role of the IL-NAshell in meth seeking

Data from our lab and others indicate that the IL-NAshell plays a nuanced role with respect to meth seeking. For example, inhibition of the IL with lidocaine had no effect on meth seeking in response to cues or drug prime^39^. In contrast, temporary inhibition of the IL with GABAa/GABAb agonists reduced cued reinstatement^20^. Neither method of pharmacological inactivation in the NAshell had an impact on drug seeking^20,39^. These disparate findings with discrete brain site manipulations underscore the importance of pathway-specific manipulations like those used herein to examine the function of neural *circuits* in particular. While our pathway manipulation is consistent with the discrete brain site inactivation of IL described in the aforementioned study^20^, we do not know how our manipulation alters output from the downstream NAshell. Although this projection is predominantly excitatory, long-range inhibitory projections from the prefrontal cortex to the nucleus accumbens have been reported^49^. It is thus possible that chemogenetic inhibition of the IL may result in no net-change in NAshell activity at the population level, but may impact activity and information processing within distinct NAshell microcircuits. It will be important to recapitulate our findings using complementary approaches to conclusively determine whether the NAshell (or NAcore for PL manipulations) is the downstream actuator of these chemogenetic effects, and to rule out non-specific effects of CNO^50^.

Notably, pharmacological inhibition of IL decreases cued heroin seeking, but NAshell inhibition produces no effect on cued reinstatement, while decreasing heroin-primed reinstatement^21^. Other, more lasting forms of IL inactivation, and “disconnection” studies of the IL-NAshell circuit, have supported a “driver” role for this circuit in heroin seeking^51,52^. Thus, our finding that the NAshell is an output by which IL activity promotes cued relapse to meth seeking is novel and suggests that there may be two or more subpopulations of neurons within the NAshell modulating relapse, since global inactivation of the NAshell only produced a marginal effect on this form of meth seeking. Indeed, there are two functionally distinct subclasses of medium spiny neurons (MSNs) in the NA: dopamine D1 receptor-expressing (D1-MSNs) and D2 receptor-expressing (D2-MSNs). Whereas evidence is lacking on the precise roles of these two subclasses of NA MSNs with regards to meth seeking, pharmacological blockade of either or both D1 and D2 receptors in the NAcore (but not the NAshell) reduces meth seeking after voluntary abstinence^59^. Evidence from cocaine and opioid models suggest that D1-MSNs promote, and D2-MSNs oppose, drug-seeking behaviors and/or behavioral sensitivity to these drugs^53–57^; but see^58^. Thus, future studies should examine whether the IL-NAshell circuit is responsible for driving cued meth relapse through NAshell D1-MSNs, which would be predicted if this same D1- vs. D2-MSN functionality holds true for meth.

Notably, our findings on the role of the IL-NAshell in cued relapse to meth seeking stand in stark contrast with the role of this circuit for cocaine. Pharmacological inactivation of the IL-NAshell circuit induces reinstatement of cocaine seeking in the absence of any relapse triggers (e.g. cues or drug prime)^13^. Further, chemogenetic activation of this circuit prior to a cued reinstatement test decreases cocaine seeking^16^. When we examined chemogenetic activation of this pathway prior to cued reinstatement of meth seeking, we found no effect. Collectively, along with our finding that chemogenetic inhibition of the IL-NAshell decreases cued reinstatement of meth seeking, these results demonstrate a functional distinction in the role of this pathway for meth vs cocaine seeking. Namely, IL projections to the NAshell promote meth seeking in the presence of meth-associated cues. The recruitment of the IL-NAshell circuit is thought to be dependent on extinction training, as chemogenetic activation of this circuit does not alter cued relapse to cocaine seeking after forced abstinence^16^. Given that all meth self-administering rats extinguished drug seeking normally over the course of extinction training, this raises the question as to which circuits might be recruited to inhibit meth seeking, or if extinction of meth seeking induces plasticity more akin to memory erasure than new learning^59,60^.

### Concluding Remarks

Using a circuit-specific chemogenetic strategy, we herein identified the functional roles of the IL and PL mPFC to NA subcircuits in meth seeking. Our results are consistent with the notion of the PL-NAcore circuit being an important projection perpetuating relapse, as inhibition of this dorsal subcircuit decreased cued relapse to meth seeking under multiple conditions, including after long-access exposure to meth and after extinction versus forced abstinence. This relapse-driving circuit may be specific to drugs of abuse, as its inhibition did not impact sucrose seeking. Furthermore, the ventral IL-NAshell subcircuit also functions to drive cued relapse to meth seeking. Thus, the mPFC to NA circuits controlling meth seeking more closely resemble those reported for heroin than cocaine.

## Acknowledgments

We thank Stewart Cox, Jordan Carter, Sammy Woods for help with surgeries, conducting the behavioral experiments, and image acquisition.

## Author Contributions

All authors discussed the experimental design, results, and interpretation of findings. AMK, JLH, and RMW conducted the experiments including the surgeries, self-administration behavior and immunohistochemistry. JLP and MDS supervised the choice and administration of viral constructs and antibodies. MDS supervised and BMS conducted the confocal microscopy. CRM conducted the statistical analysis. CMR, AMK, and JLP wrote the initial version of the manuscript and all authors edited the initial version. CMR wrote the final version of the manuscript and JLP and MDS edited the final manuscript version.

