## Supplement for "Chemogenetic Inhibition of Corticostriatal Circuits Reduces Cued Reinstatement of Methamphetamine Seeking"

**Number of Supplemental Tables:** 1
**Number of Supplemental Figures:** 6

#### Supplemental Information
#### Methods

##### 2.1 Subjects

Age-matched male (250-275g) and female (225-250g) Sprague Dawley rats (Envigo, Indianapolis, IN, USA) were used in these experiments. Rats were individually housed in a temperature and humidity-controlled vivarium on a reverse 12:12 light-dark cycle. Experiments were conducted during the rats' dark cycle. Rats were given 5 days to acclimate after arrival before treatment. Water was available ad libitum throughout the study and 20-25 g of rat chow (Envigo, Indianapolis, IN, USA) was provided daily until extinction, after which time food was provided ad libitum. All experimental procedures were approved by the Institutional Animal Care and Use Committee of the Medical University of South Carolina and were in accordance with the ‘‘Guide for the Care and Use of Laboratory Rats’’ of the Institute of Laboratory Animal Resources on Life Sciences, National Research Council.

##### 2.2 Catheter Implantation Surgery

Rats were anesthetized with ketamine (66 mg/kg i.p.; Vedco Inc, St. Joseph, MO, USA), xylazine (1.3 mg/kg i.p.; Lloyd Laboratories, Shenandoah, IA, USA), and Equithesin (0.5 ml/kg i.p.; sodium pentobarbital 4 mg/kg, chloral hydrate 17 mg/kg, 21.3 mg/kg magnesium sulfate heptahydrate dissolved in 44% propylene glycol, 10% ethanol solution) or isoflurane vaporized for inhalation (4-5% for induction in a chamber, 2%–3% through a nosecone for preparation and 1%-3% for surgical anesthesia maintenance). Ketorolac (2.0 mg/kg, i.p.; Sigma Chemical, St. Louis, MO, USA) and cefazolin (0.2 g/kg, s.c.; Patterson Veterinary, Saint Paul, MN, USA) were given before surgery as an analgesic and antibiotic, respectively. Catheters (constructed with Silastic tubing, Dow Corning Corporation, Midland, MI, USA) were inserted 4 cm into the right jugular vein and secured with silk sutures. The opposite end of the tubing ran subcutaneously and exited through a small incision on the back below the shoulder blades where an external port was exposed. The rats were flushed with 0.05 ml of TCS catheter lock solution (Access Technologies, USA) post-surgery to prevent clots and microbial growth in the catheter. Meth self-administration began at least 5 days after recovery from surgery.

##### 2.4 Meth and Sucrose Self-administration

Rats self-administered methamphetamine hydrochloride (Sigma, St Louis, MO) dissolved in sterile saline at 20mg/50ml for males and 17.5mg/50ml for females: the infusion volume was

50μl per infusion. Standard operant chambers (30 x 20 x 20 $cm^3$, Med Associates, St Albans, VT) were housed inside sound-attenuating cubicles containing a house light, a fan, two retractable levers, a drug-delivery arm attached to a swivel, and a spring leash that enclosed the tubing for drug delivery. Tygon tubing was connected to a 10 ml meth syringe fitted to an infusion pump. Fans provided white noise and ventilation. At the beginning of each session, the house light turned on and levers extended, signaling meth availability. Responding on the active lever delivered a 2s infusion of meth followed by an un-signaled 20 s timeout period where responding was without consequence. A white stimulus light positioned above the active lever and a tone signaled each meth infusion. Responding on the inactive lever was without consequence. At the end of the program, the house light turned off and levers retracted. Self-administration started 5 days after surgery. Sessions were conducted in daily 2-hour sessions beginning with a fixed ratio (FR) FR1 for 5 sessions, then advanced to FR3 for 3 sessions, and finally maintained on an FR5 for 7 sessions. The response requirement to increase the FR value was set at a minimum of 10 daily infusions. An FR5 was selected in order to clearly define lever discrimination in both sexes[26] and promote responding for cue-induced reinstatement. A subset of rats self-administered meth or sucrose (45 mg pellet; BioServ) in 8-hr daily sessions on an FR1 for 15 days and then tested for cued reinstatement following extinction.

**2.5 Extinction and abstinence**

Extinction was conducted in the self-administration chamber for at least 10 daily 2 hr sessions. Responding on either lever was without scheduled consequence. The criterion for testing was set at <25 active lever presses for two consecutive days. Abstinent rats remained in their home cage for 7 days without extinction.

**2.6 Immunohistochemistry**

At the end of the experiments, rats were transcardially perfused with phosphate buffered saline (PBS) and 10% buffered formalin, and brains removed. Tissue was sliced on a cryostat and underwent immunohistochemical procedures before being mounted onto slides, dehydrated, cover- slipped, and examined under a microscope to visualize viral expression. Subjects were eliminated from the final dataset if no expression was visible in the cell body region or if there was spread into adjacent regions defined by coronal slices from the atlas of Paxinos and Watson (2007). To visualize viral expression and validate CNO activation of DREADDs, sections were blocked with 2% normal goat serum in PBS. Sections were incubated over-night at 4°C in primary antibody: mouse Anti-Enhanced Blue Fluorescent

Protein (1:1000, Abcam: ab32791; RRID:AB_873781), chicken anti-mCherry (1:3000; LifeSpan Biosciences: #LS-C204825; RRID:AB_2716245), rabbit anti-Green Fluorescent Protein (1:1000, Abcam: ab72600; RRID:AB_1523175), and/or rabbit Fos (1:500, Millipore-Sigma:ABE457; RRID:AB_2631318). Secondary antibodies were goat anti-chicken Alexa-594 (1:1000, Invitrogen: A32759; RRID:AB_2762829), goat anti-rabbit Alexa-647 (1:1000, Invitrogen: A27040; RRID:AB_10371940), and/or goat anti-mouse Alexa-488 (1:1000, Abcam: ab150113; RRID:AB_2576208).

We determined the surgical placement of the viral constructs by visualizing the signal resulting from the viral fluorophore and comparing it to the corresponding coronal atlas plates of the medial prefrontal cortex and nucleus accumbens from Paxinos and Watson (2007). For PL, any spread into IL would result in exclusion from the data set. Some spread into the dACC was considered acceptable. For IL, which is only spans about 1mm in the DV axis, minor spread into ventral PL was considered acceptable, as ventral PL has been suggested to function similarly to IL. PL spans approximately 3mm in the DV axis, and any spread beyond the most ventral 1mm portion would have resulted in exclusion for the IL target.

##### 2.7 Confocal and Microscopy and Analysis

Briefly, Alexa-594 amplified mCherry and Alexa-647 amplified Fos were imaged using a Leica SP8 laser-scanning confocal microscope (Leica microsystems, Wetzlar, Germany) equipped with HyD detectors for enhanced sensitivity. mCherry was imaged using an OPSL 552nm laser line and Fos was imaged using a Diode 638nm laser line using a 20X air objective (0.75 N.A.). Pinhole, Diode 638 laser power, and gain were empirically determined prior to the experiment and were then held constant. OPSL 552 laser power was held relatively constant; adjusting only to avoid saturation. Z-stacks were acquired at a frame size of 1024x1024, 0.5 um Z-step size, with 2-line averages. Z-stacks were ~20 µm in the Z-plane. 3-6 sections were imaged per animal in either the PL or IL cortices. Each image consisted of one hemisphere.

Deconvolved (Huygens, SVI, Netherlands) Z-stacks were exported to Imaris (v. 9.0). A constant baseline subtraction was then applied to remove background for the Fos channel. The spots tool was used to semi-automatically count the number of mCherry+ cells (20 µm diameter) and Fos+ nuclei (15 µm diameter). The spots colocalization MATLAB extension was then used to detect mCherry+/Fos+, mCherry+/Fos-, and mCherry-/Fos+ cells (5 µm distance between spots maximum). Falsely recognized, or obviously missed overlapping and non-overlapping cells were removed or added, respectively, by the investigator. The number of mCherry+Fos+/mCherry neurons was calculated and exported. Data was collapsed across hemispheres and across sections, then expressed as an animal average.

**Supplemental Figure 1.**

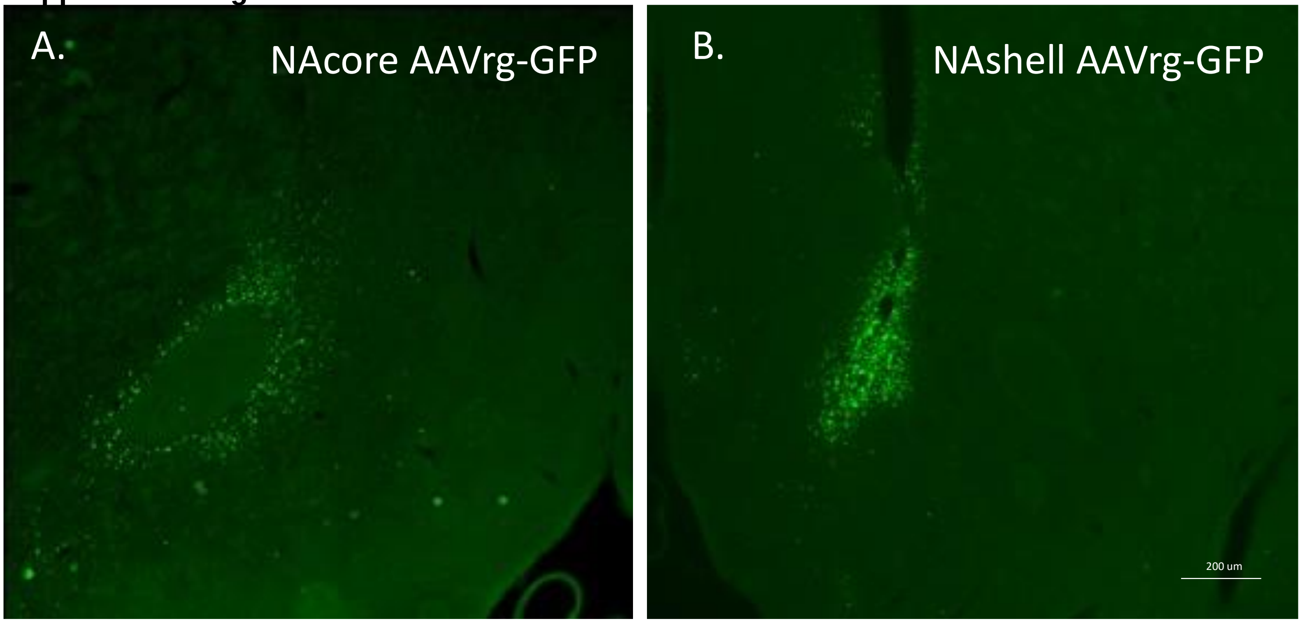

**Supplemental Figure 1.** Representative images of the terminal locations infused with AAVrg-pmSyn1-Cre-GFP (AAVrg-Cre). **A)** Image depicts green florescent protein expression (GFP) in the NAcore (stereotaxic coordinates: AP +1.6 mm; ML ± 2.8 mm (10° angle); DV -7.1 mm. **B)** Image depicts GFP in the NAshell (stereotaxic coordinates: AP +1.7 mm; ML, ± 0.8 mm, DV -7.0 mm). Combined these images represent the viral spread in the terminal areas showing restriction to the core vs. shell. Images were acquired at 2.5 mm.

#### Supplemental Figure 2.

Timeline and experimental descriptions

Arrival Acclimation → Surgery → Recovery → Self-administration → Extinction Abstinence → Testing → Viral placement

| Experiment | Virus Infusions | | Meth | | | | |
|---|---|---|---|---|---|---|---|
| | PFC | NA | Self-administration | Extinction | Tests | N's | Analysis |
| 1. Inhibition of PL-NAcore | AAV-DIO-hM4Di | AAVrg-Cre | 2 hr day FR 1, 3, 5 | 2 hr 10 days | 2 hr Veh, CNO (3 or 10 mg/kg) | 23 | Mixed effects |
| 2. Inhibition of PL-NAcore | AAV-DIO-hM4Di | AAVrg-Cre | 2 hr day FR 1, 3, 5 | abstinence | 1 hr Veh, 10 mg/kg CNO | 19 | Unpaired t |
| 3. Activation of PL-NAcore | AAV-DIO-hM3Dq | AAVrg-Cre | 2 hr day FR 1, 3, 5 | 2 hr 10 days | 2 hr Veh, 10 mg/kg CNO | 16 | Mixed effects |
| 4. CNO control | AAV-DIO-mCherry | AAVrg-Cre | 2 hr day FR 1, 3, 5 | 2 hr 10 days | 2 hr Veh, CNO (3 or 10 mg/kg) | 21 | Mixed effects |
| 5. Inhibition of PL-NAcore | AAV-DIO-hM4Di. | AAVrg-Cre | 8 hr day FR 1 | 2 hr 8 days | 2 hr Veh, 10 mg/kg CNO | 8 | Repeated measures |
| 6. Inhibition of PL-NAcore | AAV-DIO-hM4Di. | AAVrg-Cre | Sucrose 8 hr day FR 1 | 2 hr 8 days | 2 hr Veh, 10 mg/kg CNO | 8 | Repeated measures |
| 7. Inhibition of IL-NAshell | AAV-DIO-hM4Di. | AAVrg-Cre | 2 hr day FR 1, 3, 5 | 2 hr 10 days | 2 hr Veh, CNO (3 or 10 mg/kg) | 12 | Mixed effects |
| 8. Activation of IL-NAshell | AAV-DIO-hM3Dq | AAVrg-Cre | 2 hr day FR 1, 3, 5 | 2 hr 10 days | 2 hr Veh, 10 mg/kg CNO | 16 | Mixed effects |

**Supplemental Figure 2.**

Rats in all experiments arrived at MUSC and were given at least 5 days to acclimate to the new environment. They underwent jugular catheterization and intracranial viral infusions. After recovery, rats went through self-administration, extinction or abstinence and cued reinstatement of meth seeking. Finally, rats were euthanized and viral placements were evaluated. Overall, eight experiments were conducted. There specific methods are described in the table. [PL, prelimbic; IL, Infalimbic; NA, nucleus accumbens; Veh, vehicle (dimethylsulfoxide, DMSO); CNO, clozapine-N-oxide; FR, fixed ratio].

**Supplemental Figure 3.**

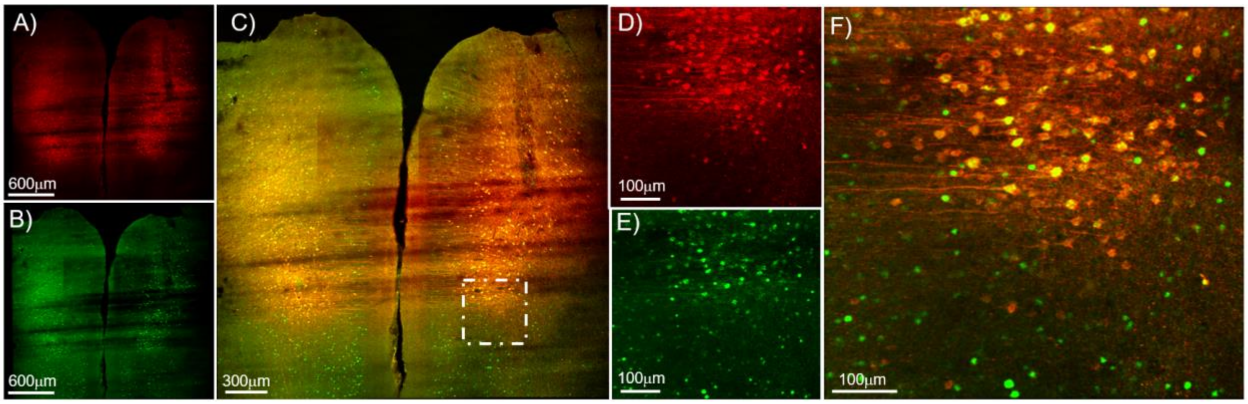

**Supplemental Figure 3.** Representative images of Fos+/mCherry+ expression in the prelimbic (PL) regions of the prefrontal cortex. **A)** Low magnification image from a rat that had AAV-hSyn-DIO-hM4D(Gi)-mCherry [0.6 µL/side at 0.15µL/min] infused into the PL and AAVrg-pmSyn1-Cre (AAVrg-Cre) into the Nucleus Accumbens (NA) core. DREADD expressing cells are shown in red **B)** Immunohistochemical detection of Fos (green) in the same section. **C)** Merged image of DREADD-mCherry expression (red) and Fos (green). This rat was tested under vehicle (dimethylsulfoxide, DMSO) conditions in the presence of methamphetamine associated cues and tissue was collected following a 2 hr test session. As such, the Fos seen here is indicative of neuronal activity induced by methamphetamine paired cues. **D)** Higher magnification images are shown of the inset region (hatched box) in panel C. Here as before DREADD-mCherry cells are shown in red. **E)** Corresponding Fos expression in the inset region is shown in green. F) The merged image of the inset region is shown. This figure depicts mCherry+ DREADD expression localized in PL, which was present mostly in layers V and II/III. Additionally, the final panel demonstrates that as expected, there are some additional Fos+ cells outside of the PL.

#### Supplemental Figure 4.

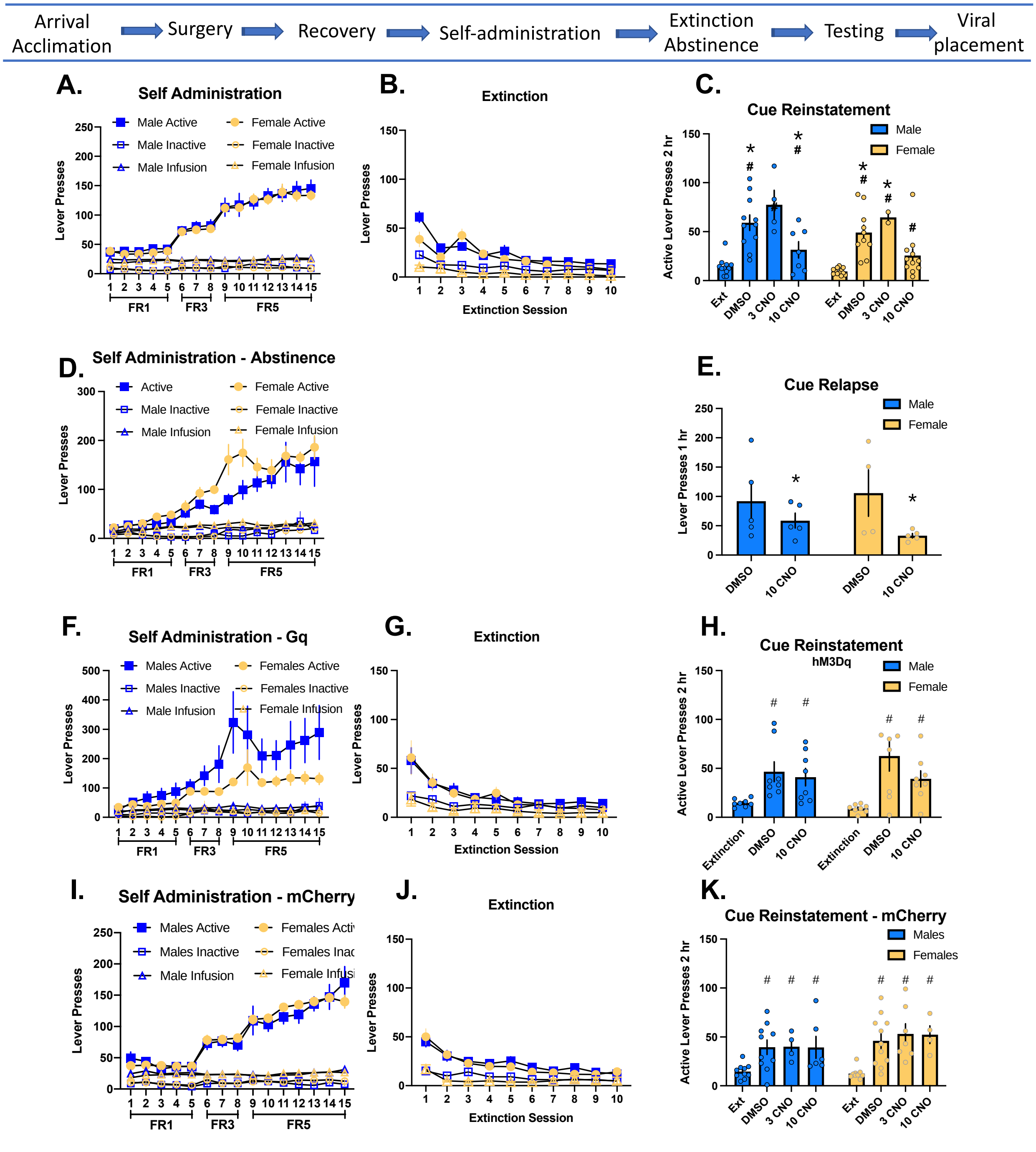

**Supplemental Figure 4.** Self-administration (SA), extinction, and reinstatement sessions disaggregated by sex for rats that had prelimbic (PL)-nucleus accumbens (NA) core Cre-dependent circuit manipulation. SA and extinction sessions were 2 hr. Rats tested in the presence of methamphetamine associated cues and test sessions were 1-2 hr as depicted on the Y-axis. (A-C) Self-administration, infusions, extinction, and reinstatement following PL-NAcore DIO-hM4Di inhibition in males and female rats. Inhibition occurred through ip injection of clozapine-N-oxide (CNO, 3 and 10 mg/kg) or dimethylsulfoxide (DMSO) was used as a vehicle/control condition. (D-E) Self-administration and cued responding following PL-

NAcore DIO-hM4Di inhibition in male and female rats that underwent abstinence (*indicates main effect of test condition). (F-G) Self-administration, infusions, extinction, and meth cued reinstatement in male and female rats that had AAVrg-mCherry infused into the PL-NAcore. (I-J) Self-administration, infusions, extinction, and reinstatement of PL-NAcore DIO-hM3Dq activation in male and female rats. Activation occurred through ip injection of CNO 10 mg/kg or DMSO was used as a vehicle/control condition. Figure SF1A, D, F, and I are combined in the main text as Figure 2A. Figure SF1B, G, and J are combined in Figure 2B. Figure SF1C, E, H, and K are depicted in Figure 3 (main text) collapsed across sex.
*Significant difference from 10 mg/kg CNO.
#Significant difference from extinction.

**Supplemental Figure 5.**

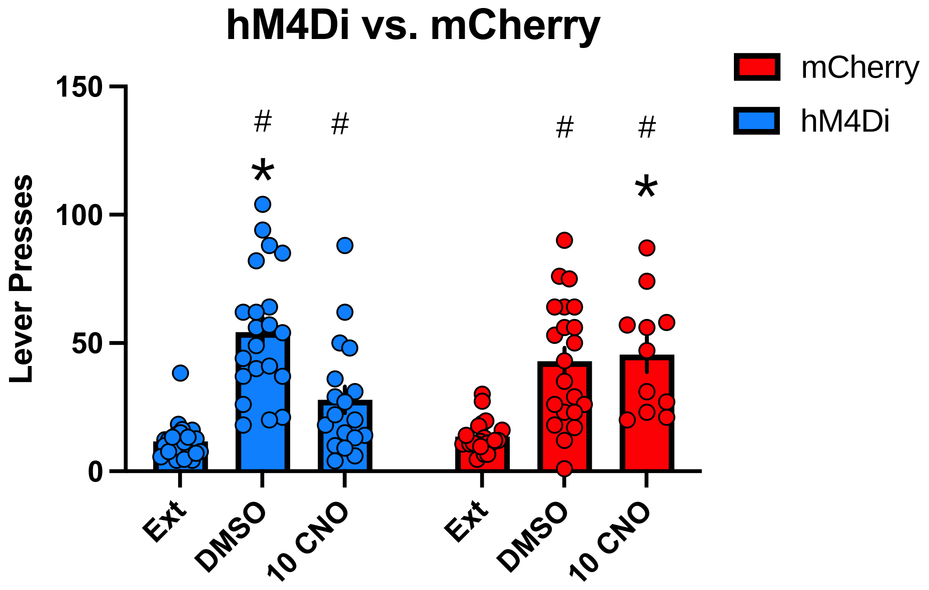

**Supplemental Figure 5.** Direct comparison between rats that had DIO hM4Di or DIO mCherry infused into the prelimbic (PL) cortex reveal a significant interaction between DIO viral infusion and test day [$F(2,67)=5.37$, $p<0.007$]. Specifically, hM4Di rats injected with dimethylsulfoxide (DMSO) or clozapine-N-oxide (CNO) had increased lever responding in response to methamphetamine (meth) associated cues. Whereas, cue reinstatement was significantly reduced in hM4Di rats injected with CNO relative to DMSO. (Holme-Sidak's, $p<0.05$). Rats with DIO-mCherry in the PL reinstated responding to meth associated cues regardless of whether they were injected with DMSO or CNO (Holme-Sidak's, $p<0.05$). In fact, CNO increased responding in the viral mCherry control group above that of the DIO DREADD inhibition with hM4Di. This pattern demonstrates that our effects of PL-NAcore inhibition is specific to CNO inhibition and not non-specific viral infusion or non-specific effects of CNO.
* Significant difference from hM4Di 10 CNO, $p<0.05$
### Significant difference from extinction.

#### Supplemental Figure 6.

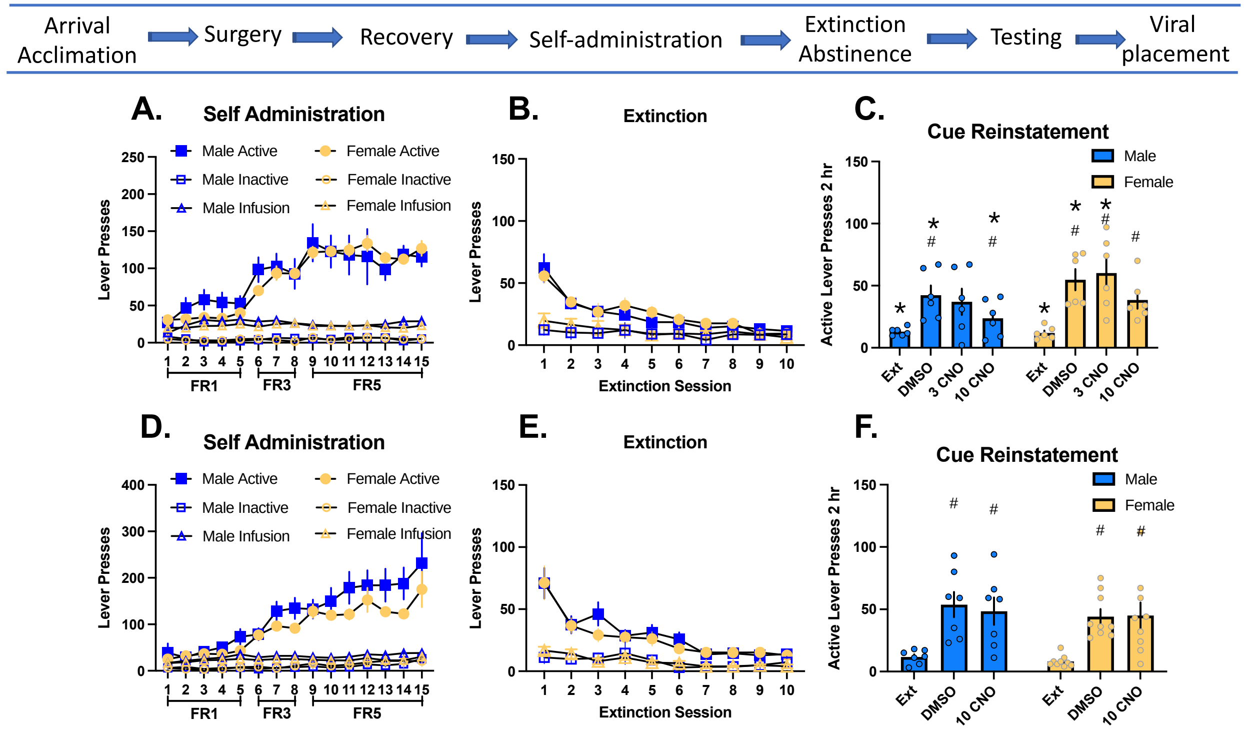

**Supplemental Figure 6.** Self-administration, extinction, and reinstatement sessions disaggregated by sex for rats that had Infralimbic (IL)-nucleus accumbens (NA) shell Cre-dependent circuit manipulation. (A-C) Self-administration, infusions, extinction, and reinstatement following IL-NAshell DIO-hM4Di inhibition in males and females. Inhibition occurred through ip injection of clozapine-N-oxide (CNO, 3 and 10 mg/kg) or dimethylsulfoxide (DMSO) was used as a vehicle/control condition. (D-F) Self-administration, infusions, extinction, and reinstatement of IL-NAshell mCherry male and female control rats. Activation occurred through ip injection of CNO 10 mg/kg or DMSO was used as a vehicle/control condition. SF4A and D are combined in the main text as Figure 5A and SF4B and E are combined in Figure 5B. Figure SF4C and F are depicted in Figure 5 collapsed across sex in the main text.

*Significant difference from 10 mg/kg CNO.

#Significant difference from extinction.

**Supplemental Table 1.**

Supplemental Table 1. Inactive lever responding on reinstatement tests.

**PL-NAcore**

| Experiment | Prelimbic | Accumbens Core | VEHICLE (DMSO) | 3 CNO | 10 CNO |
|---|---|---|---|---|---|
| Reinstatement | hM4Di | rAAV | 5.53 (±1.32) | 7.86 (±1.9) | 2.81 (±0.6) |
| Relapse | hM4Di | rAAV | 12.9 (±2.34) | | 10.8 (±2.23) |
| Reinstatement | hM3Dq | rAAV | 7.56 (±1.83) | | 10.25 (±2.6) |
| Reinstatement | mCherrry | rAAV | 5.39 (±1.31) | 7.44 (±3.14) | 7.11 (±3.14) |
| Long Access meth | hM4Di | rAAV | 9.12 (±1.27) | | 48.00 (±2.14) |
| Long Access sucrose | hM4Di | rAAV | 4.29 (±0.86) | | 6.7 (±2.01) |

**IL-NAshell**

| Experiment | Infralimbic | Accumbens Shell | VEHICLE (DMSO) | 3 CNO | 10 CNO |
|---|---|---|---|---|---|
| Reinstatement | hM4Di | rAAV | 5.33 (±1.26) | 5.67 (±1.47) | 7.15 (±1.39) |
| Reinstatement | hM3Dq | rAAV | 4.14 (±1.27) | | 4.4 (±1.24) |

**Supplemental Table 1** provides the means and standard errors for inactive lever responding on the test session. Inactive lever responding during self-administration and extinction are depicted in the main text figures and are divided in the supplement by sex.
